# The Structured Coalescent and its Approximations

**DOI:** 10.1101/091058

**Authors:** Nicola F. Müller, David A. Rasmussen, Tanja Stadler

## Abstract

Phylogenetics can be used to elucidate the movement of genes between populations of organisms, using phylogeographic methods. This has been widely done to quantify pathogen movement between different host populations, the migration history of humans, and the geographic spread of languages or the gene flow between species using the location or state of samples alongside sequence data. Phylogenies therefore offer insights into migration processes not available from classic epidemiological or occurrence data alone. Phylogeographic methods have however several known shortcomings. In particular, one of the most widely used methods treats migration the same as mutation, and therefore does not incorporate information about population demography. This may lead to severe biases in estimated migration rates for datasets where sampling is biased across populations. The structured coalescent on the other hand allows us to coherently model the migration and coalescent process, but current implementations struggle with complex datasets due to the need to infer ancestral migration histories. Thus, approximations to the structured coalescent, which integrate over all ancestral migration histories, have been developed. However, the validity and robustness of these approximations remain unclear. We present an exact numerical solution to the structured coalescent that does not require the inference of migration histories. While this solution is computationally unfeasible for large datasets, it clarifies the assumptions of previously developed approximate methods and allows us to provide an improved approximation to the structured coalescent. We have implemented these methods in BEAST2, and we show how these methods compare under different scenarios.

## 1 Introduction

The relatedness of samples of homologous genetic sequences assuming asexual reproduction are the result of a past branching process. The same applies to other sources of data, such as languages or phenotypic markers. This past branching process contains information about ancestral population histories and can be inferred from data using phylogenetic trees. One of the information encoded in such trees is the structure of a population and the movement of information (e.g. genes or words) between subpopulations. *Phylogeographic* methods allow to elucidate such information given the state or location of samples. Phylogeographic methods have been used to analyze the global spread of influenza viruses (Bedford et al., 2010; Bahl et al., 2011; Lemey et al., 2014; Bedford et al., 2015), the origins of HIV-1 (Faria et al., 2014) and various other diseases (Bourhy et al., 2008; Raghwani et al., 2011). Analogously to the analysis of epidemics, such methods have been used to study the geographic origin of brown and polar bears (Edwards et al., 2011). Related methods have been used to study the evolutionary history of humans (Gronau et al., 2011) and great apes (Mailund et al., 2012). The same methods have also been applied to study the origin of the Indo-European language family (Bouckaert et al., 2012).

A range of phylogeographic methods for inferring population structure from phylogenies have been proposed. The mugration method (Lemey et al., 2009) treats migration as a continuous time Markov chain, such as used to model mutation, and assumes the migration process to be independent of the tree generating process. This assumption can lead to biases in estimates of migration rates when sampling is biased (De Maio et al., 2015). Other methods, such as those based on the structured coalescent (Takahata, 1988; Hudson, 1990; Notohara, 1990) and the related isolation-with-migration models (Wakeley, 2000; Nielsen and Wakeley, 2001; Hey, 2010), do not make this independence assumption. In contrast to the mugration-based methods, they require the state (or location) of any ancestral lineage in the phylogeny at any time to be inferred (Beerli and Felsenstein, 2001; Ewing et al., 2004; Vaughan et al., 2014). Inferring lineage states is computationally expensive, as it normally requires Markov chain Monte Carlo (MCMC) based sampling, and limits the complexity of scenarios that can be analyzed.

Other approaches (Volz, 2012; Palczewski and Beerli, 2013) seek to marginalize over all possible migration histories by treating lineage states probabilistically instead of using MCMC based sampling. Rather than assigning lineages to particular states, the probability of each lineage being in each state is calculated at all times using a set of previously described differential equations (Volz, 2012). Such a marginalization approach (rather than explicit sampling of states) allows for the analysis of larger datasets (De Maio et al., 2015). While this approach appears to only make the assumption of lineage independence, i.e. that the state or location of one lineage does not depend on any other lineage (De Maio et al., 2015), it remains unclear if there are additional assumptions not being accounted for.

In this paper, we derive an exact numerical solution of the structured coalescent with discrete states for neutrally evolving asexual populations. This solution is based on the joint probabilities of lineages being in any possible configuration. However, it quickly becomes computationally unfeasible for more than a few lineages and states. It allows us however to clarify the assumptions used in previous approaches (Volz, 2012; De Maio et al., 2015) and to develop a more refined approximation to the structured coalescent. We then show how the different approximations compare in terms of tree, parameter and root state inference under both biased and unbiased sampling conditions. Simulations reveal that our new approximation outperforms previous approximations at comparable computational cost. We then apply these different approximations to a previously described avian influenza virus dataset (Lu et al., 2014) sampled from different regions of North America to show that the choice of method influences the interpretation of data in practice.

## 2 New Approaches

Approaches that calculate the probability density of a phylogeny under the structured coalescent given a set of coalescent and migration rates typically use MCMC to integrate over possible migration histories. Using this Monte Carlo integration however heavily limits the size of datasets that can be analyzed. Already at a small number of different states, efficiently exploring the space of all possible migration histories becomes unfeasible. Methods that are able to integrate over these migration histories but avoid MCMC sampling hold great promise in their ability to analyse larger datasets. We therefore derive an exact solution to the structured coalescent process with discrete states for neutrally evolving asexual populations that integrates over all possible migration histories using ordinary differential equations. We refer to this approach as ESCO, the exact structured coalescent.

While ESCO is exact, it requires solving a number of differential equations that is proportional to the “number of different states” to the power of the “number of co-existing lineages”. This originates from the need to calculate the probability of every possible configurations of a set of co-existing lineages and states using migration and coalescent rates. We therefore develop a lower-dimensional approximation that is based on keeping track of the marginal lineage state probabilities instead. We call this approach the marginal lineage states approximation of the structured coalescent (MASCO). This approach allows us to reduce the number of differential equations that have to be solved between events to “number of states” times “number of lineages”, but ignores any correlations between lineages. Using this approach, the state of a lineage is calculated backwards through time, integrating over potential migration events and incorporating the probability of no coalescences between branching events in the phylogeny. This means that the state or location of a lineage is directly dependent on the coalescent process. In particular, the observation that two lineages that do not coalesce for a longer time are unlikely to be in the same state is incorporated in this approach.

In comparison to MASCO, we show that the approach of (Volz, 2012) requires the additional assumption that the state of a lineage evolves independently of the coalescent process between events. This means that changes in the probabilities of lineages being in a certain state are only dependent on the migration rates, and are completely independent of other lineages in the phylogeny. We refer to this approach as SISCO, the state independence approximation of the structured coalescent. The differential equations describing how lineages evolve between events for ESCO and MASCO are both derived in the *Materials and Methods* section. Whereas the differential equations for SISCO have been derived previously (Volz, 2012).

## 3 Results

### 3.1 Tree height distributions under the structured coalescent and its approximations

The structured coalescent and its approximations describe different probability distributions over trees. To see how these distributions compare, we performed direct backwards-in-time simulations under the structured coalescent using MASTER (Vaughan and Drummond, 2013), analogously to Vaughan et al. (2014). These trees were compared to trees sampled under ESCO, MASCO, SISCO, as well as BASTA (De Maio et al., 2015), a numerical approximation of SISCO. Under these latter four models, trees were sampled from their respective probability distributions using MCMC in BEAST2 (Bouckaert et al., 2014). Since it is difficult to directly compare distributions of trees, we instead compared the distribution of tree heights.

For each of the five scenarios (direct, ESCO, MASCO, SISCO, BASTA) and three different overall migration rates, we obtained 8000 trees. We used a model with three different states, sampling three, two and one individuals from each state, respectively. Coalescent rates were different in each state (λ_1_ = 1, λ_2_ = 2, λ_3_ = 4) and migration rates were different between states (*m*_1,2_ = 1, *m*_1,3_ = 2, *m*_2,1_ = 0.1, *m*_2,3_ = 0.3, *m*_3,1_ = 1, *m*_3,2_ = 1). To show how the different methods perform under different overall migration rates, the rates between states were scaled with 1 (fast migration), 0.1 (medium migration) and 0.01 (slow migration).

Figure S1 shows the distribution of tree heights sampled using MCMC and compares them to the distribution of tree heights obtained by directly simulating trees under the structured coalescent. Of the different methods, only the distribution of ESCO is consistent with direct simulation. Only keeping track of the marginal lineage states (MASCO) leads to slightly shorter tree heights. Further assuming lineage states to be independent of the coalescent process (SISCO) results in greatly underestimated tree heights. BASTA (De Maio et al., 2015), being an approximation of SISCO, performs very similar to SISCO. The shorter tree heights under SISCO compared to MASCO can be explained in the following way. Not taking into account how the coalescent process influences lineage states leads to an overestimation of the probability of two lineages being in the same state if no coalescent event is observed by SISCO compared to MASCO. Overestimating the probability of two lineages being in the same state then also leads to a higher probability of them coalescing. This in turn results in shorter trees since lineages are expected to coalesce at a faster rate. SISCO and BASTA in general perform worse at slower migration rates than at rates in the same order of magnitude as the rates of coalescence.

### 3.2 Root state probabilities

The ancestral state or location of lineages back in time is often of interest for biological questions. For example, in a pathogen phylogeny the root location is informative of the geographic origin of an epidemic. Here we show on one fixed tree how the exact structured coalescent compares in the inference of the root state to its approximations. We additionally inferred the root state using MultiTypeTree (Vaughan et al., 2014), which uses MCMC to sample lineage states and does not rely on approximations, to obtain a reference root state probability (Vaughan et al., 2014). We inferred the probability of the root being in either state for different migration rates in one direction while holding the rate in the other direction constant.

The exact structured coalescent and the one only keeping track of the marginal lineage states (MASCO) agree well with the inferred posterior mean using MultiTypeTree (Figure S2). The inferred state probabilities using SISCO on the other hand do not, showing that the assumption of independence between the lineage states and the coalescent process does not only describe a wrong probability distribution over trees but can also leads to biased inference of ancestral states.

### 3.3 Estimation of migration rates

Coalescent methods are often used to infer population and migration parameters from trees. To show how the inference of the migration rates compares to the true rate, we simulated 1000 trees under the structured coalescent with symmetric migration rates from 10^−5^ to 1 and pairwise coalescent rates of 2 using MASTER. Hence, we consider a range of cases from very strong to very weak population structure, where the probability of migration is in the same order as coalescence. Each tree consisted of 4 contemporaneously sampled leafs from each of the two states. We fixed the coalescent rates to the truth, assumed symmetric migration rates and then inferred the maximum likelihood estimate of the migration rate using the exact structured coalescent (ESCO) and its approximations MASCO and SISCO.

The results are summarized in Figure S3. When only keeping track of the marginal lineage states (MASCO), the migration rates are estimated well. Making the further assumption of independence of the lineage states and the coalescent process (SISCO) leads to strong biases in estimates of the migration rates. The lower the migration rates are compared to the coalescent rates, the greater the underestimation of the migration rates becomes.

### 3.4 Estimation of rate asymmetries

In the previous section, we inferred the rate of migration given (or conditional on) the true coalescent rate and the information that the migration rates were the same in both directions. In reality, these rates can greatly vary across states or locations. It is therefore important for methods to be able to perform well in situations where rates are asymmetric. Previous work showed that the ability to infer migration rate asymmetries greatly depends on the method used (De Maio et al., 2015). Here we compare inferences of rate asymmetries under MASCO and SISCO. Applying ESCO to the same trees would not be computationally feasible, due to the larger number of lineages existing in parallel.

Figure S4 shows the median ratios of inferred coalescent and migration rates using MASCO and SISCO. The estimates of coalescent rate ratios (Figure S4 top row) are accurate under both simulation scenarios and methods. Estimates of the migration rate ratios are biased in the presence of asymmetric coalescent rates (Figure S4 bottom left) using SISCO, but not MASCO. SISCO overestimates the backwards in time migration out of the state with a faster coalescent rates and into the state with a slower coalescent rate. An underestimation of the rate in the other direction was observed as well. When the coalescent rates are symmetric, both methods are unable to capture very strong asymmeries in the migration rate ratios (Figure S4 bottom right). However, when taking into account the highest posterior density (HPD) intervals of the estimates, most estimates contain the true rate ratio (see figures S1 and S2). MASCO is overall better at infering those migration rate asymmetries than SISCO.

### 3.5 Sampling bias

Previous work showed that the approximate structured coalescent is able to accurately infer migration rates even when sampling fractions are biased, given samples are taken contemporaneously (De Maio et al., 2015). Here we explore the effect of biased sampling fractions in the presence of serial sampling. We compare the exact structured coalescent ESCO to its approximations MASCO and SISCO.

Figure S5 reveals that ESCO is able to unbiasedly infer the migration rates in both directions, independent of sampling biases or migration rates. The same applies to MASCO. For SISCO however, biased sampling leads to an underestimation of the backwards migration rate into the oversampled state and an overestimation of the rates into the undersampled state for intermediate and high migration rates. At low migration rates, both rates are underestimated.

### 3.6 Application to Avian Influenza Virus

To show how the inference of the origin of an epidemic varies with the method used, we applied the two approximations of the structured coalescent (MASCO and SISCO) to a previously described avian influenza dataset (Lu et al., 2014; De Maio et al., 2015) to infer the geographic location of the root.

In Figure S6, we show the inferred region of the root using MASCO and SISCO. Despite the fact that almost all samples from the central US were collected after 2009 and that samples from the East Coast and the North West fall closer to the root, SISCO places the root with over 80% probability in the central US. MASCO on the other hand places the root to be most likely at the East Coast, one of the least likely root locations according to SISCO. Also, in contrast to SISCO it doesn’t exclude most regions from being the location of the root based on the phylogenetic data available. We provide a possible explanation to why we observe differences in the inferred root state in the Discussion below.

## 4 Discussion

We provide an exact way to calculate the probability density of a phylogenetic tree under the structured coalescent (Takahata, 1988; Hudson, 1990; Notohara, 1990) without the need to sample migration histories, as in previously described approaches (Beerli and Felsenstein, 2001; Ewing et al., 2004; Vaughan et al., 2014), by solving a set of ordinary differential equations.

Additionally, we introduce a new approximation that outperforms a previously described approximation (Volz, 2012). This new approximation facilitates a trade-off between speed and accuracy. The increased speed compared to the exact solution originates from ignoring any correlations between lineages. This assumption leads to better scaling of the computational complexity with the number of states and lineages. We show that this assumption allows us to infer migration, coalescent rates and root states in all scenarios tested within this simulation study.

Additionally assuming independence of the lineages states from the coalescent process, as introduced in (Volz, 2012), leads however to major biases in parameter and root state inference. These biases are especially pronounced in our simulations when migration is slow compared to the coalescent rate. This observation can be explained in the following way:

The lower migration rates are compared to coalescent rates, the stronger the influence of the coalescent process on the configuration of lineages across states. The assumption of independence of the lineage states from the coalescent process does not allow for the incorporation of this information into the calculation of lineage state probabilities though.

Next, we showed how the approximations of the structured coalescent perform in inferring asymmetric coalescent and migration rates. While coalescent rates are inferred accurately for both approximations, inference of migration rate ratios is biased when coalescent rates are asymmetric under SISCO. We also showed that under biased sampling, inferences of migration rates are strongly biased under SISCO, but not under MASCO.

Both biases can be understood in the following way. A lineage may have a higher probability of coalescing in one state than another either because the pairwise coalescent rate in one state is higher (e.g. due to a smaller effective population size) or because more lineages reside in one state than another (e.g. because of biased sampling). Taking the influence of the coalescent process on lineage states into account, as done under MASCO, reduces the probability of a lineage occupying a state with a high coalescent rate if no coalescent events occur.

In other words, MASCO redistributes the probability mass assigned to each state to reflect the observed coalescent history, including the observation that a lineage may have not yet coalesced (see equation 3). SISCO does not redistribute probability mass to reflect the observation that a lineage has not yet coalesced. In order to reduce the probability of lineages coalescing in a state with high rates of coalescence, it overestimates the migration rate out of such states. This overestimation of migration rates out of a state is observable when having asymmetric coalescent rates due to either a higher pairwise coalescent rate within a state or having more lineages in a given state due to biased sampling. Either way, the migration rate out of the state with a higher coalescent rate is overestimated and underestimated in the other direction. While revising this manuscript, it was brought to our attention that updates to the R package rcolgem (Erik M Volz, 2016) (based on Volz (2012)), uses a related approach to redistribute probability mass between states.

While MASCO does redistribute probability mass via the coalescent process, it ignores the correlations between lineages encoded in the joint probabilities when only considering marginal lineage state probabilities. These correlations are expected to be especially strong in parts of the tree where there are only a few co-existing lineages present. These correlations between lineages are induced by the coalescent process. The rate at which lineages coalesce is highly dependent on the number of lineages in a state. Having one or two lineages in the same state is the difference between having a zero or non-zero rate of coalescence, whereas having a 1000 or a 1001 lineages in the same state doesn’t impact the rate of coalescence as much. In turn, this means that at lower number of lineages, the state of a single lineage has a much larger impact on the rate at which coalescent events are expected. This then leads to stronger correlations between the state of individual lineages. We however did not find a scenario under which MASCO would be considerably biased compared to the exact description of the structured coalescent.

We applied the different approximations of the structured coalescent to avian influenza virus HA sequences sampled from different orders of birds in North America. We found that the inferred region of the root varies with the method used. SISCO places high confidence in the center of the USA being the root state. MASCO on the other hand infers the East coast to be the most likely location of the root, while also placing a considerable amount of probability mass to other locations such as the North East or North West to reflect uncertainty in the phylogenetic data about the root location.

Asymmetric coalescent rates may offer one explanation why SISCO places more probability on the center being the root location than MASCO and why it excludes all other states from being possible root states. We have shown that asymmetric coalescent rates can bias the inference of migration rates. Under SISCO, asymmetric coalescent rates lead to an overestimation of the migration rate from a state with fast coalescent rate into a state with slow coalescent rate and an underestimation of the migration rates in the other direction (recall that we consider backwards in time rates). Because the coalescent rate in the center is inferred to be low, SISCO puts much more weight on it being the source than MASCO. The opposite appears to occur for the East Coast, which is inferred to have a very high rate of coalescence. MASCO infers the East Coast to be the most likely source region while it is almost excluded using SISCO.

Although we used the AIV analysis to illustrate how inferences obtained from MASCO and SISCO can differ, the results presented here should be interpreted with caution with regards to any biological implications as we ignored population structure arising between different avian host species. We additionally assumed coalescent and migration rates to be constant over time, potentially further biasing the inference of the root state.

While population dynamics such as changing transmission (i.e. coalescent) and migration rates through time can greatly influence the shape of a phylogeny, we ignored such dynamics in this study. However, compared to mugration type methods (Lemey et al., 2009), the structured coalescent approximation introduced here can be extended in a conceptually straightforward way to allow for dynamic populations (Volz et al., 2009; Volz, 2012). The improved approximation to the structured coalescent introduced here should therefore allow for more accurate quantification of pathogen movement in structured populations with complex population dynamics while still being computationally efficient enough to be applied to large datasets. Lastly, while we do not consider the special case of isolation-with migration (Wakeley, 2000; Nielsen and Wakeley, 2001; Hey, 2010), the here presented approaches should translate easily by i) assuming contemporenous sampling (which is already possible) and ii) combining states after times *t*_1_, …, *t*_*m* − 1_, allowing the combined state to have new coalescent and migration rates.

## 5 Materials and Methods

### 5.1 Principle of the structured coalescent process

The structured coalescent (Takahata, 1988; Hudson, 1990; Notohara, 1990) extends the standard coalescent by allowing lineages to occupy different states. If we consider *L_i_* to be a random variable that denotes the state of lineage *i* with state space {1, …, *m*}, there are *m^n^* different possible configurations 𝒦 of how *n* lineages can be arranged (𝒦 = (*L*_1_ = *l*_1_, …, *L_i_* = *l_i_*, …, *L_n_* = *l_n_*), *l_i_* ∈ {1, …, *m*}). These configurations can change over time by adding and removing lineages or by lineages changing state. Throughout this paper, we consider time going backwards from present to past, as typically done under the coalescent.

A migration event along one lineage *i* from state *a* to state *b* changes the configuration of lineages as follows:

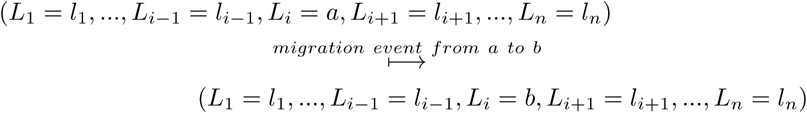

In figure S7, this corresponds to lineage 1 in *blue* changing to *red*.

Configurations can additionally change due to sampling. Sampling events simply add lineages, such as *L*_3_ = *red* is added in figure S7. Typically, we condition on the sampling events, but one can also introduce a rate for samples being obtained.

A coalescent event between lineage *i* and *j* with *i* < *j* changes the configuration as follows:

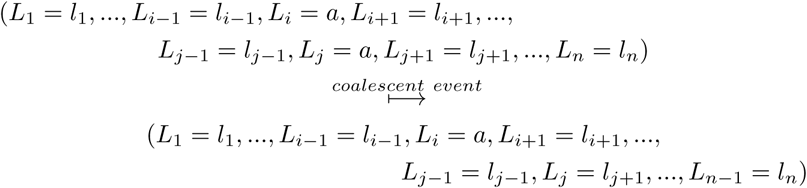

Lineages *j* + 1, ˙., *n* are relabelled to *j*, …, *n* − 1 and lineage *i* denotes the parent lineage of *i* and *j* after a coalescent event. The most recent coalescent event in figure S7 for example changes the configuration from (*L*_1_ = blue, *L*_2_ = blue, *L*_*3*_ = *red*) to (*L*_1_ = *blue, L*_2_ = *red*).

The rate at which coalescent events in state *a* happen can be calculated from the pairwise coalescent rate λ_*a*_ in state *a* and the number of lineages *k_a_*(𝒦) in state *a* for a given configuration 𝒦. The pairwise coalescent rate denotes the rate at which any two lineages in a state coalesce. For a given configuration 𝒦, the total rate 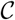 at which coalescent events between any two lineages in the same state happen is:

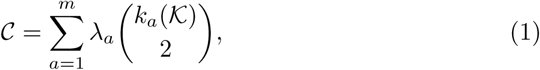

where 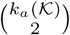 is the number of pairs of lineages in state *a* given configuration 𝒦. Under the standard Wright-Fisher model, the pairwise coalescent rates, λ_*a*_, are the inverse of the effective population sizes *N*_*e_a_*_.

### 5.2 Calculating the likelihood for a tree under the structured coalescent

Structured coalescent methods typically use MCMC to integrate over possible lineage state configurations along a tree (Beerli and Felsenstein, 2001; Ewing et al., 2004; Vaughan et al., 2014). This is sometimes referred to as sampling migration histories. Given a migration history, the likelihood for a tree can be calculated under the structured coalescent with given migration and coalescent rates. Here, we want to calculate the marginal likelihood for a tree without sampling those migration histories, but by integrating over all possible migration histories *H*. Formally, we seek to calculate the following probability:

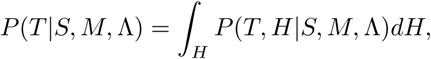

with *T* being the tree, *S* the sampling states of the tips, *M* the set of migration rates and Λ the set of coalescent rates.

Let *P*_*t*_(*L*_1_ = *l*_1_, …, *L*_*i*_ = *l*_*i*_, …, *L*_*n*_ = *l*_*n*_*, T*) be the probability density that the samples more recent than time *t* evolved according to the coalescent history, i.e. the branching pattern, given by our tree *T* between the present time 0 and time *t* and that the *n* lineages at time *t, L*_1_, …, *L_n_*, are in states *l*_1_, …, *l_n_*. In figure S7, this probability is the joint probability of a configuration at time *t* with the lineages being either in red or blue, and the probability of the branching pattern and tip states being as observed between time *t* and 0 (ignoring the particular configurations in that time interval).

We aim to calculate *P_t_* for *t* = *t_mrca_*, with *t_mrca_* being the time of the root of the tree *T*. At the root of the tree, summing over the probability of the remaining lineage being in any state will yield the likelihood for the tree, 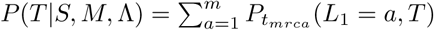.

In order to evaluate *P_t_* at *t* = *t_mrca_*, we start at the time of the most recent sample, at *t* = 0, and iteratively calculate *P*_*t*+Δ*t*_ based on *P_t_*. To calculate *P_t_*, we split the calculation into three parts: time intervals in the tree where no coalescent or sampling events happen, sampling events, and coalescent events. Below, we first consider the interval part of this calculation. Afterwards, we calculate the contribution of coalescent and sampling events.

#### Interval contribution

For the interval part, we calculate *P*_*t*+Δ*t*_ based on *P_t_* allowing for no event in time step Δ*t* (second line below), observing a migration event leading to the configuration at *t* + Δ*t* (third line below), or seeing more than one event (i.e. higher order terms which are of order *O*((Δ*t*)^2^) leading to the configuration at *t* + Δ*t* (forth line below):

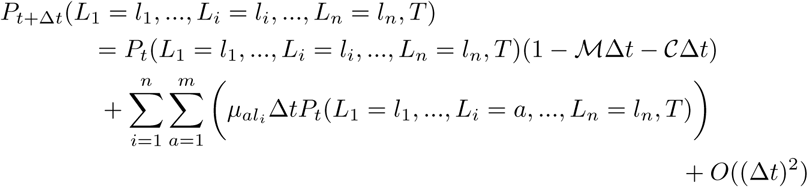

Here, ℳ is the sum of migration rates and 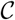 the sum of coalescent rates for configuration (*L*_1_ = *l*_1_, …, *L_i_* = *l_i_*, …, *L_n_* = *l_n_*). The rate *μ_al_i__* denotes the rate at which migration events from *a* to *l_i_* happen. Now, when re-arranging and letting Δ*t* → 0, we obtain the differential equation,

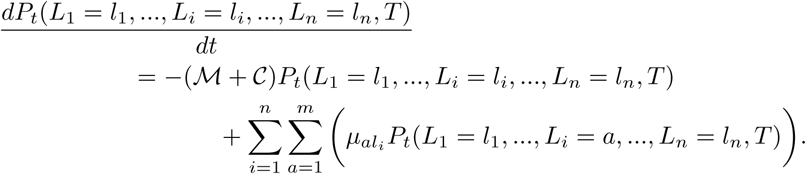

With explicitly writing ℳ and 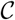 (using equation 1 for 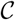), we obtain,

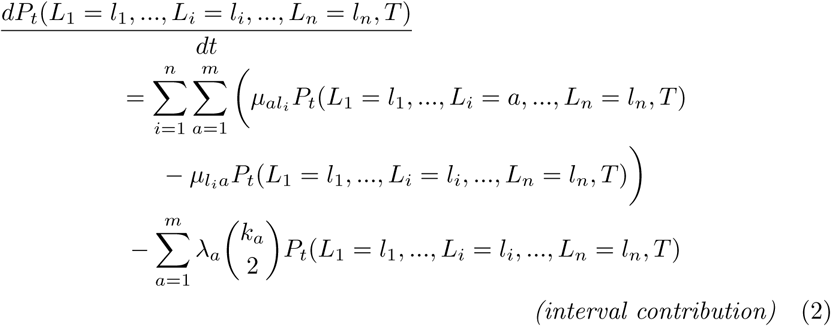

with the double summation on the right hand side considering the contribution of migration and the fourth line considering the contribution of coalescence. Further, 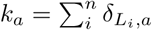 where *δ* is the Kronecker delta with *δ*_*L_i_,a*_ = 1 for *L*_*i*_ = *a* and 0 otherwise. Note that in the case of *l*_*i*_ = *a*, the two terms in the migration part cancel each other out and the net migration is 0. This *interval contribution* equation allows us to calculate *P*_*t*_ within intervals by solving the differential equation.

It is important to note that this differential equation shows a direct link between the coalescent process and the probability of a set of lineages being in a configuration. For example, configurations that would favor high coalescent rates among lineages would become less probable over intervals during which no coalescent events occur in the tree.

#### Sampling event contribution

At every sampling event the state of the sampled lineage is independent of all other lineages in the tree. We can therefore calculate the probability of any configuration at a samping event at time *t* as follows:

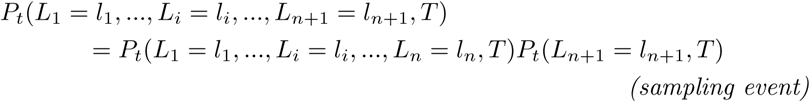

In scenarios where the sampling state is known to be say *a*, we have *P*_*t*_(*L*_*n*+1_ = *a,T*) = 1 and *P*_*t*_(*L*_*n*+1_ = *b,T*) = 0 for *b* ≠ *a*. In cases where the sampling state is an inferable parameter or not exactly known, this probability can be between 0 or 1.

#### Coalescent event contribution

Next, we have to calculate the probability of the new configuration resulting from a coalescent event between lineages *i* and *j* in state *a* at time *t*. This probability can be expressed by the following equation:

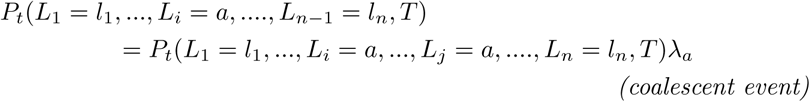

Thus based on the three equations, *(interval contribution), (sampling event), (coalescent event)*, we can calculate the likelihood for a tree, *P*(*T*|*S*,*M*, Λ). We refer to this approach as the exact structured coalescent (ESCO).

### 5.3 Approximations of the exact structured coalescent

Between events (sampling and coalescent), the exact structured coalescent requires *m*^*n*^ differential equations to be solved, with *m* being the number of different states and *n* the number of lineages present at a point in time. To be able to analyse datasets with more than a few states and lineages, approximations have to be deployed.

In the exact structured coalescent, the state of a lineage *i* is always associated with a configuration 𝒦 and the coalescent history described by the tree *T*. Keeping track of these configurations automatically keeps track of all correlations between lineages. We will now assume that lineages *i, j* and *k* and their states *l*_*i*_, *l*_*j*_ and *l*_*k*_ are uncorrelated, i.e.:

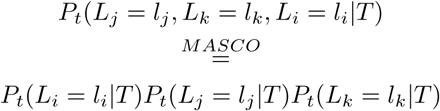

Using this approximation, we will write down an expression for:

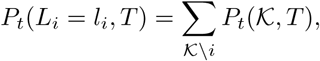

with 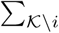 being the summation over all configurations while fixing the state of lineage *i*.

The interval contribution, i.e. the change in marginal lineage state probability over time, 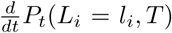, can be derived from equation 2 and the above expression employing the MASCO assumption. This derivation is explained step by step in the supplement, and results in the following differential equation:

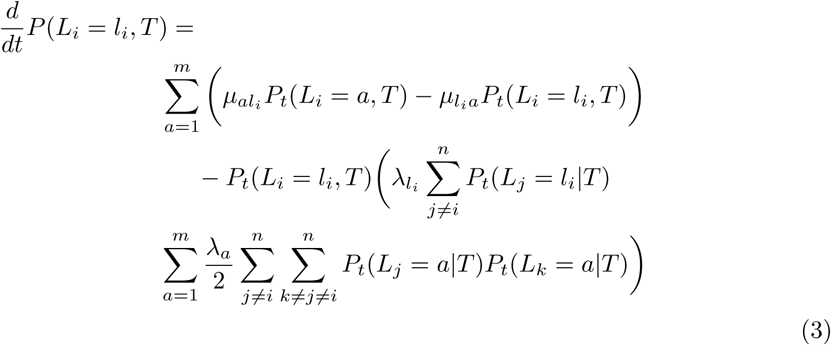

The second line denotes the change in marginal lineage state probability due to migration. The third line denotes the reduction in *P*(*L*_*i*_ = *l*_*i*_, *T*) due to the rate of coalescent events directly involving lineage *i*. The fourth line now denotes the rate of coalescence of events that do not involve lineage *i*. Integrating equation 3 over time is equivalent to calculating the probability that the lineage *i* is in state *l*_*j*_ and that all lineages evolved up to time *t* as given by the coalescent history *T*. The above equation ensures that 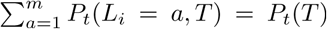 for every lineage *i*. This is due to the fact that *P*_*t*_(*L*_*i*_ = *a, T*) incorporates information about everything that happens in the tree and therefore also events not directly involving lineage *i*.

For the coalescent event contribution, we calculate the probability of lineage *i* coalescing with lineage *j* in state *a* as,

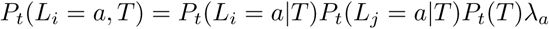

with 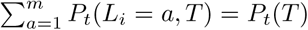 being the probability of having observed the coalescent history *T* up to time *t*, and *P*_*t*_(*L*_*i*_ = *a*|*T*) = *P*_*t*_(*L*_*i*_ = *a, T*)/*P*_*t*_(*T*) where *P*_*t*_(*L*_*i*_ = *a, T*) is obtained through Eqn. 3. As with ESCO, we relabel the indices of all lineages after each coalescent event such that the labels of n co-existing lineages is always *i* ∈ {1, …, *n*}. Note that since we keep track of the joint probabilities of lineages being in any state and the coalescent history *T*, the probabilities of all lineages k not involved in the coalescent event have to be updated as well, such that 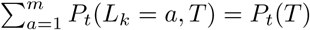 holds at every point in time for every lineage *k*.

For the sampling event contribution, we simply add a lineage *n* + 1 with associated probability *P*_*t*_(*L*_*n*+1_ = *l*_*n*+1_, *T*), such that ∑_*l*_*n*+1__ *P_t_*(*L*_*n*+1_ = *l*_*n*+1_,*T*) = *P_t_*(*T*).

The likelihood for a given tree under the structured coalescent under the MASCO approximation now is, 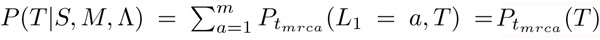.

A further approximation to the interval contribution can be obtained by ignoring the two coalescent terms in equation 3, i.e. additionally assuming independence of the lineage states from the coalescent process between events. Thus, we assume that lineages move independent of the coalescent process between events:

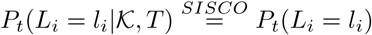

This allows to simplify equation 3 to:

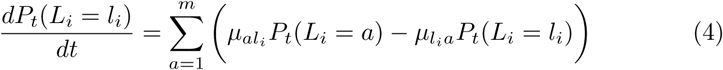

and:

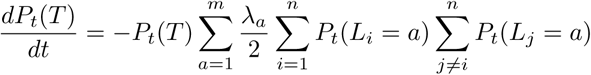

The derivation of the two equations above is explained step by step in the supplement. At a coalescent event between lineage *i* and *j*, the probability of *P_t_*(*T*) can be updated as follows:

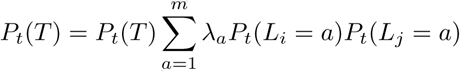

Similarly, we can calculate the probability of the parent lineage being in state *a* as:

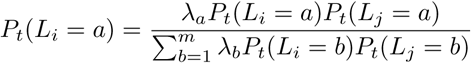

The likelihood for a given tree under the structured coalescent under the SISCO approximation now is, 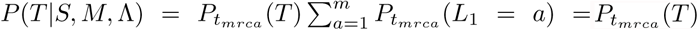.

We refer to this as the state independence approximation of the structured coalescent (SISCO). The equations used by SISCO to calculate the state of a lineage over time have been described previously in Volz (2012). While these lineage state probabilities evolve independently of the coalescent history *T* between events, they do depend on *T* at sampling and coalescent events.

### 5.4 Application to Avian Influenza Virus

We applied the different approximations of the structured coalescent to a previously described data set of Avian Influenza Virus H7 hemaglutinen (HA) sequences (Lu et al., 2014), sampled from the bird orders Anseriformes, Charadriiformes, Galliformes and Passeriforms in Canada, Mexico and the USA. We used previously aligned sequences from De Maio et al. (2015). The sequences were analyzed in BEAST2 (Bouckaert et al., 2014) using an HKY+Γ_4_ site model. A strict molecular clock model was assumed and the first two and the third codon positions were allowed to have different mutation rates. MASCO and SISCO were used as structured coalescent population priors. The dataset was split into 7 different states according to geographic regions in North America (see table S1). Three parallel MCMC chains were run for 1 * 10^7^ (MASCO) resp. 2 * 10^7^(SISCO) iterations with different initial migration and coalescent rates. After a burnin of 10%, the chains were combined and the probability of the root being in each state was assessed. The combined chain had ESS values above 100 for any inferred probability density or parameter.

### 5.5 Implementation

We implemented all three approximations in one common package for BEAST2. ESCO and MASCO use a forth order Runge-Kutta solver with fixed step size implemented in the Apache Commons Math library (http://commons.apache.org) to solve equations 2 and 3. SISCO uses matrix exponentiation to solve the lineage state probabilities over time (equation 4). All three structured coalescent methods use pairwise coalescent rates and backwards in time migration rates as described above. In the Results section, we present simulation analyses highlighting the quality of the different structured coalescent approximations.

### 5.6 Software

Simulations were performed using a backwards in time stochastic simulation algorithm of the structured coalescent process using MASTER 5.0.2 (Vaughan and Drummond, 2013) and BEAST 2.4.2 (Bouckaert et al., 2014). Script generation and post-processing were performed in Matlab R2015b. Plotting was done in R 3.2.3 using ggplot2 (Wickham, 2009). Tree plotting and tree height analyses were done using ape 3.4 (Paradis et al., 2004) and phytools 0.5-10 (Revell, 2012). Effective sample sizes for MCMC runs were calculated using coda 0.18-1 (Plummer et al., 2006).

### 5.7 Data availability

All scripts for performing the simulations and analyses presented in this paper as well as the Java source code for the structured coalescent methods are available at https://github.com/nicfel/The-Structured-Coalescent.git. Output files from these analyses, which are not on the github folder, are available upon request from the authors.

## 6 Acknowledgement

NM and TS were funded in part by a SNF SystemsX grant (TBX). DR is funded by the ETH Zürich Postdoctoral Fellowship Program and the Marie Curie Actions for People COFUND Program. TS is supported in part by the European Research Council under the Seventh Framework Programme of the European Commission (PhyPD: grant agreement number 335529).

---

**Figure S1:**
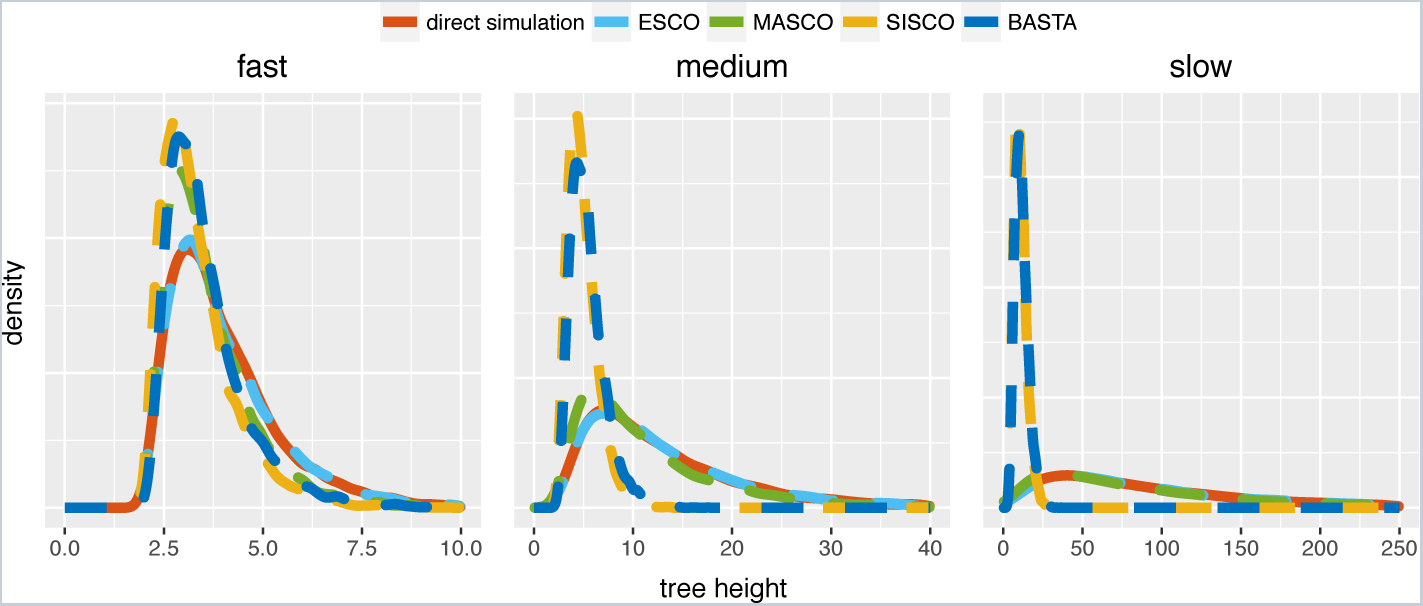
Comparison of MCMC sampled to simulated tree heights using the different structured coalescent methods. Sampled tree heights when the rates of migration are fast, i.e. in the same order of magnitude as coalescence. When the rates of migration are medium, i.e. one order of magintude lower than coalescence and slow, i.e. two orders of magnitude lower than coalescence. The trees were sampled using MCMC for one million iterations, storing every thousandth step, after a burnin of 20%.

**Figure S2:**
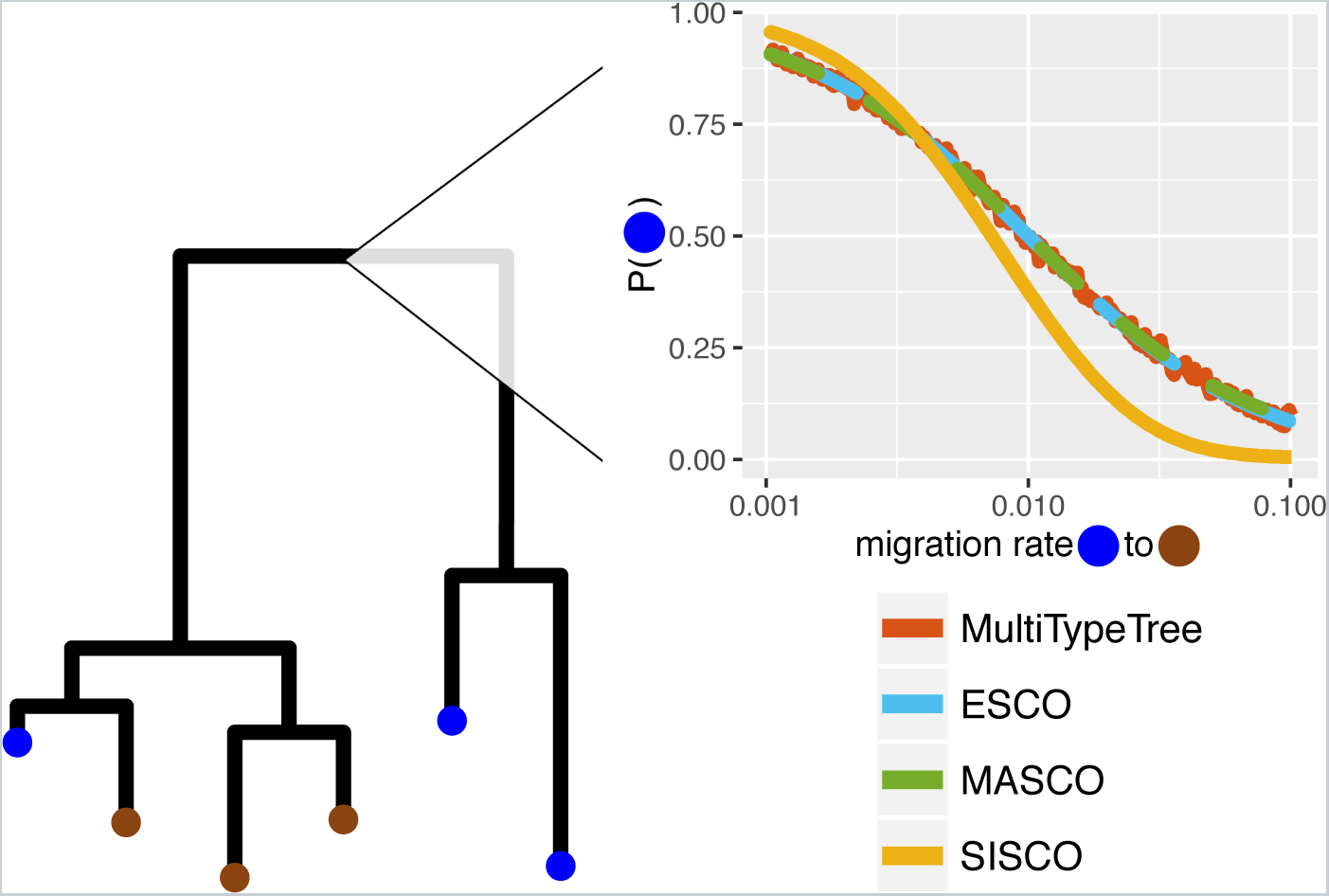
Inferred location of the root for different migration rates and structured coalescent approaches. The plot shows the probability of the root being in the blue state (y-axis) depending on the migration rate from blue to brown (x-axis), for the given tree and sampling states. The migration rate from brown to blue was held constant at 0.01.

**Figure S3:**
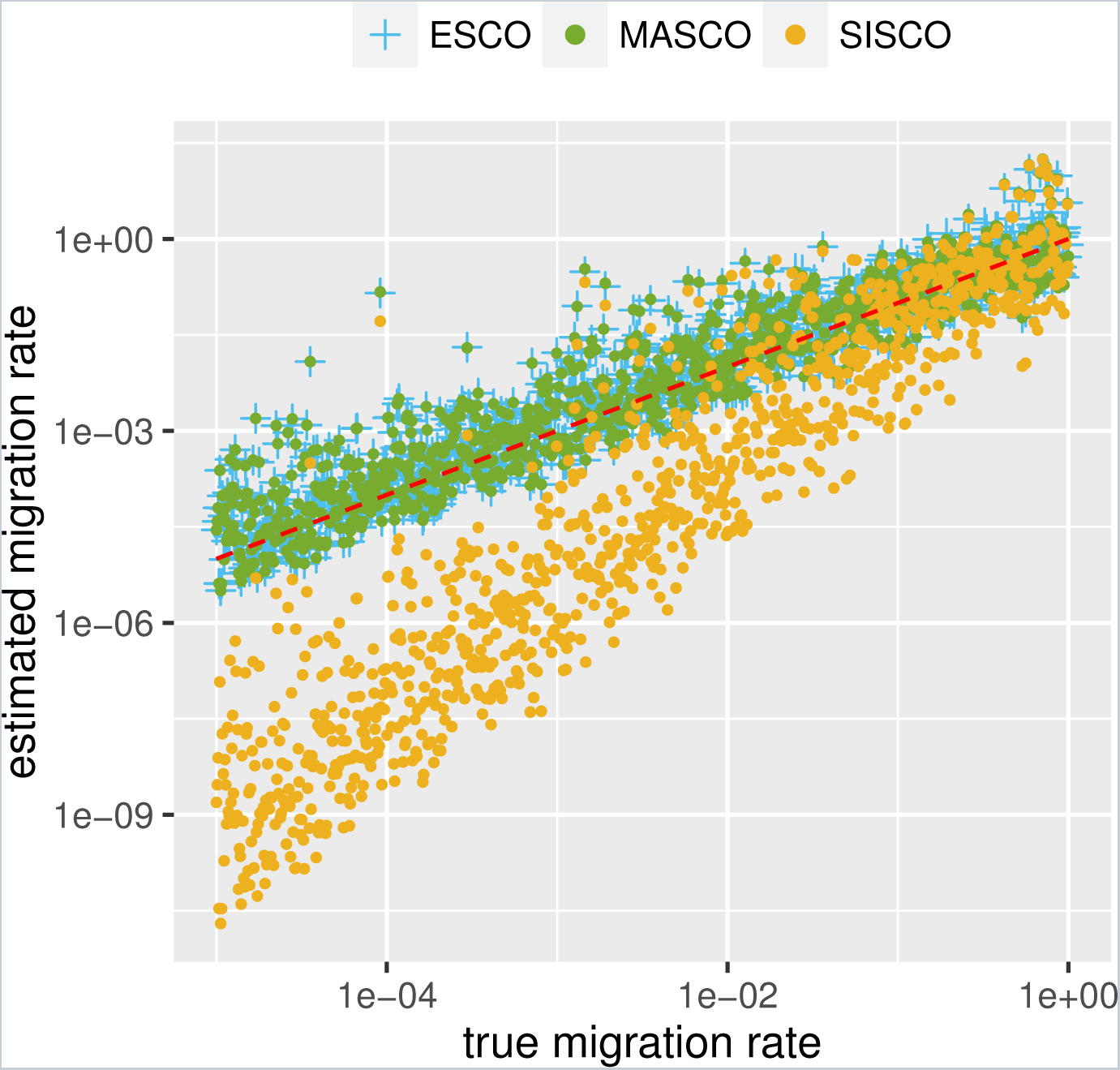
Maximum likelihood estimates of migration rates using the exact structured coalescent and its approximations. Here we compare simulated migration rates (x-axis) to the maximum likelihood estimates of the migration rate (y-axis), estimated using the exact structured coalescent ESCO and its approximations MASCO and SISCO. The coalescent rates are fixed to the truth, and the migration rates are assumed to be symmetric. The red line indicates where the true values should lie.

**Figure S4:**
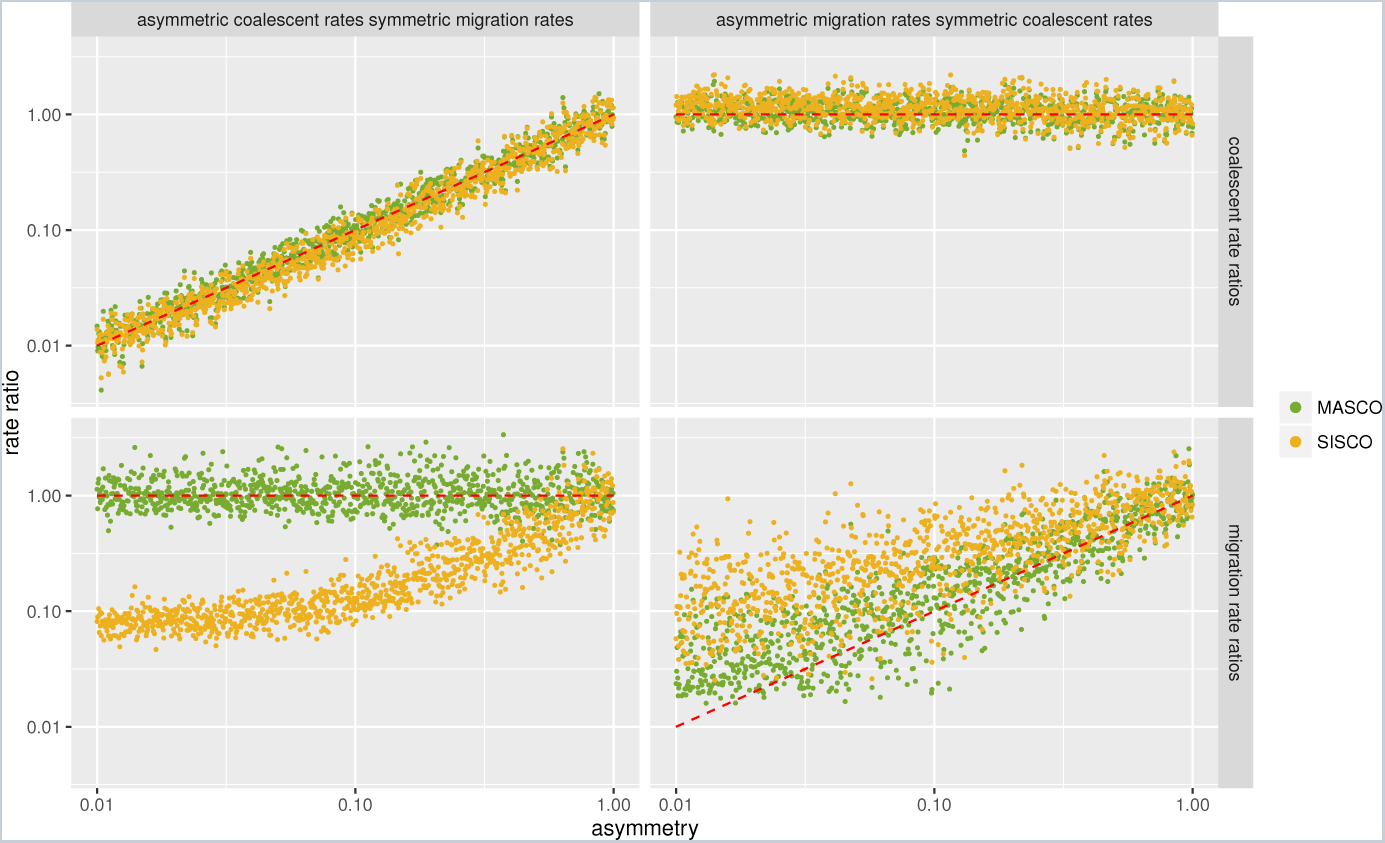
Inferred asymmetry of migration and coalescent rates. Here we show the inferred median coalescent (upper row) and migration (lower row) rate ratios under different conditions. In the first column, the coalescent rate ratios (x-axis) are varied while the migration rates ratios are kept constant. In the second column, the migration rate ratios (x-axis) are varied, while the coalescent rate ratios are kept constant. We simulated a total of 2000 trees using MASTER with 100 tips from each of the two different states sampled uniformly between times t=0 and t=10. Of these trees, 1000 were simulated with pairwise coalescent rate ratios λ_1_/λ_2_ from 0.01 to 1, λ_1_+λ_2_=4 and migration rates in both directions equal to 1. The other 1000 trees were simulated with migration rate ratios from *m*_12_/*m*_21_ from 0.01 to 1, m_12_+m_21_ = 2 and pairwise coalescent rates in both states equal to 2, using exponential priors with mean 2 for the coalescent rates and mean 1 for the migration rates Both coalescent rates and both migration rates are estimated. The red line indicates where the estimates should lie.

**Figure S5:**
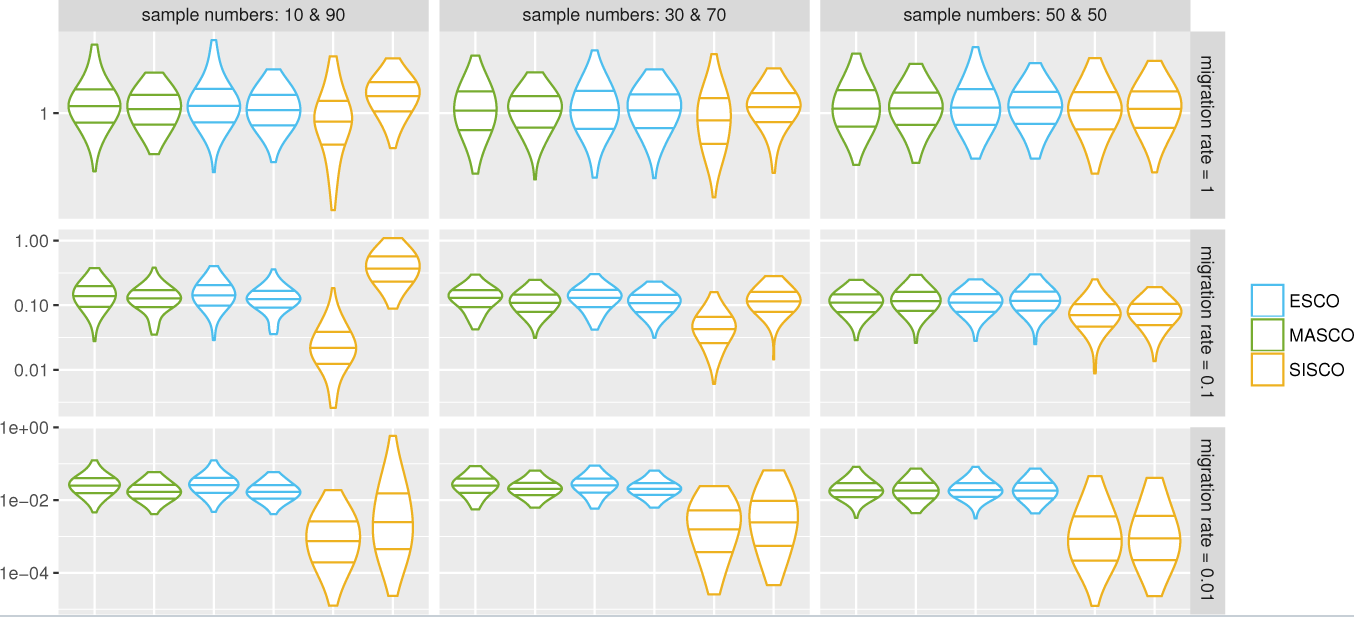
Inferred migration rates under different sampling conditions. The plot shows the distribution of mean inferred migration rates using ESCO, MASCO and SISCO. From the left, the first distribution of a color (indicating the different methods) always shows the distribution of mean inferred migration rates from state 1 to state 2. The second distribution from the same color shows the rates from state 2 to 1. From left to right the number of samples from state 1 and state 2 are changed, while from top to bottom the true symmetric migration rates are going from 1 to 0.01. The lines within the violin plots indicate the 25%, 50% and 75% quantiles. The coalescent rates were 2 in both states and the migration rates ranged from 0.01 to 1. The migration rates were always symmetric, i.e. the same in both directions. The lefs we sampled uniformly between *t* = 0 and *t* = 25. Each simulation was repeated 100 times and each inference was run with 3 parallel MCMC chains, each with different initial values. An exponential prior distribution with the mean = 1 was used on the migration and coalescent rates.

**Figure S6:**
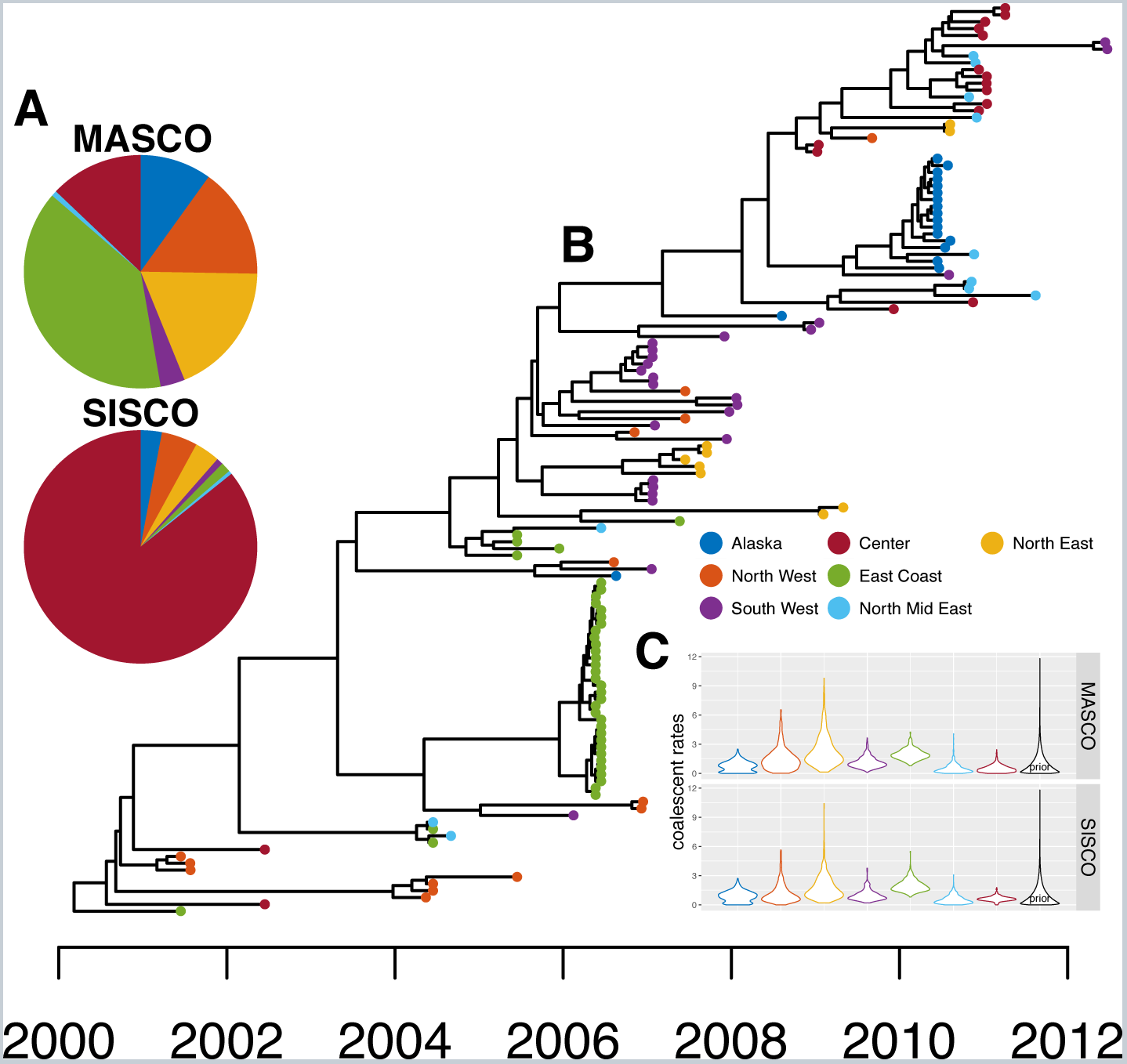
Inference of the root regions of AIV sampled from different places in North America. A Maximum clade credebility tree inferred from AIV sequences sampled in different regions of the USA, Canada and Mexico using MASCO as a population prior. The node heights represent the mean node heights. The tip color indicate the different sampling regions shown in the legend. B Inferred root regions using MASCO (top) and SISCO (bottom). The pie charts show the inferred probability of the root being in either of the different states/regions by MASCO and SISCO. C Violin plots of the inferred coalescent rates for the different regions. The black plot distribution is the exponential prior with mean 1. We used this prior for both coalescent and migration rates.

**Figure S7:**
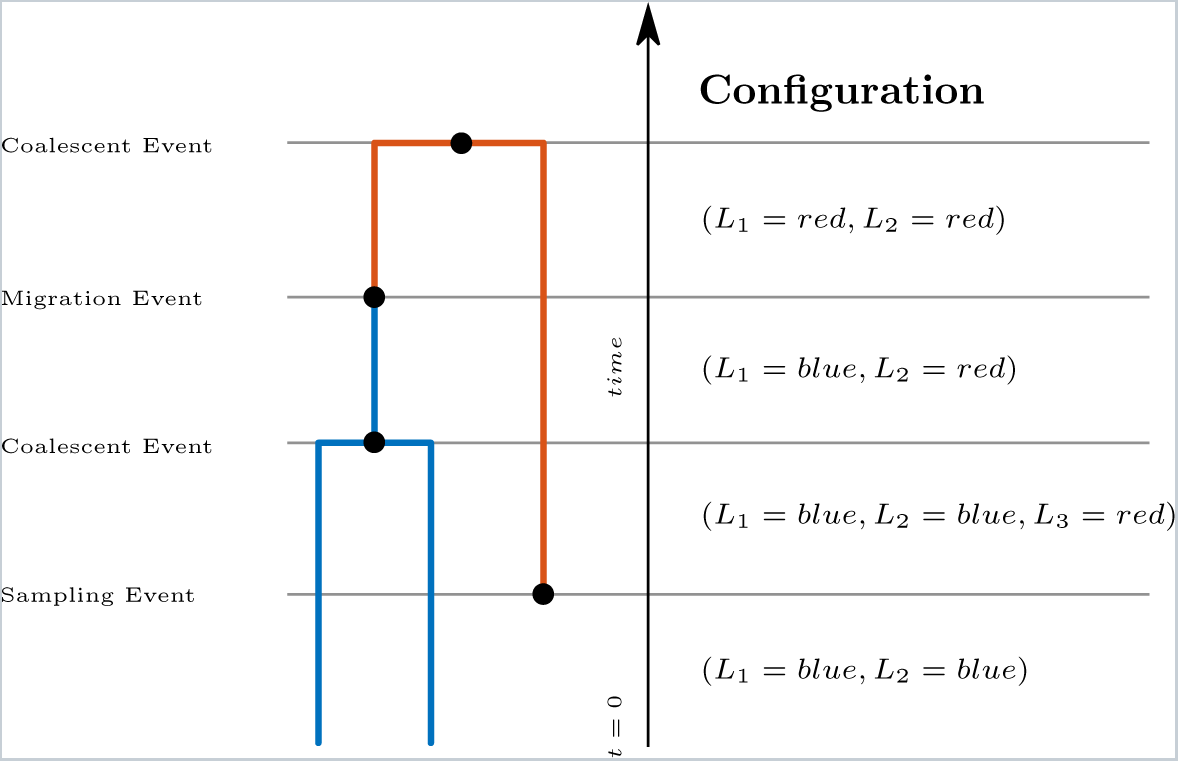
Events and configurations on an example tree. Here, we illustrate the possible events and the configurations before and after each event on a simple tree, with time going backwards from present to past. The first two lineages are both in state blue, i.e. the configuration is (*L*_1_ = *blue, L*_2_ = *blue*), with lineage 1 being the parent lineage of 1 and 2 after relabelling. After a lineage in state *red* is sampled, the configuration changes, as given in the figure. A coalescent event in state blue then reduces the number of lineages in state blue to 1. A migration event then causes lineage *L*_1_ to change state from *blue* to *red*.

